# The ash dieback invasion of Europe was founded by two individuals from a native population with huge adaptive potential

**DOI:** 10.1101/146746

**Authors:** Mark McMullan, Maryam Rafiqi, Gemy Kaithakottil, Bernardo Clavijo, Lorelei Bilham, Elizabeth Orton, Lawrence Percival-Alwyn, Ben J. Ward, Anne Edwards, Diane G.O. Saunders, Gonzalo Garcia, Jonathan Wright, Walter Verweij, Georgios Koutsovoulos, Kentaro Yoshida, Tsuyoshi Hosoya, Louisa Williamson, Philip Jennings, Renaud Ioos, Claude Husson, Ari M. Hietala, Adam Vivian-Smith, Halvor Solheim, Dan MaClean, Christine Fosker, Neil Hall, James K.M. Brown, David Swarbreck, Mark Blaxter, Allan Downie, Matthew D. Clark

## Abstract

Accelerating international trade and climate change make pathogen spread an increasing concern. *Hymenoscyphus fraxineus,* the causal agent of ash dieback is one such pathogen, moving across continents and hosts from Asian to European ash. Most European common ash *(Fraxinus excelsior)* trees are highly susceptible to *H. fraxineus* although a small minority (~5%) evidently have partial resistance to dieback. We have assembled and annotated a draft of the *H. fraxineus* genome which approaches chromosome scale. Pathogen genetic diversity across Europe, and in Japan, reveals a tight bottleneck into Europe, though a signal of adaptive diversity remains in key host interaction genes (effectors). We find that the European population was founded by two divergent haploid individuals. Divergence between these haplotypes represents the 'shadow' of a large source population and subsequent introduction would greatly increase adaptive potential and the pathogen's threat. Thus, EU wide biological security measures remain an important part of the strategy to manage this disease.

---

Human movement and climate change may increase the incidence of plant pathogen introductions as new environments become favourable^1^. Dutch elm disease is one example of an emerging pathogen epidemic which caused the loss of billions of elm trees globally^2^. More recently, ash dieback caused by the ascomycete fungus, *Hymenoscyphus fraxineus,* is causing severe dieback symptoms and death in European common ash *Fraxinus excelsior* and *F. angustifolia* (narrow-leaved ash). *H. fraxineus* jumped host from ash species native to Asia to *F. excelsior*^3,4^. In its native range, *H. fraxineus* is a leaf pathogen^5^ with little impact on its host, whereas in Europe it is killing European ash at an alarming rate and displacing the nonaggressive indigenous fungus *H. albidus*^6,7^. The likelihood that a pathogen can infect two plant species increases with their phylogenetic relatedness^8^ and so, host jump epidemics like ash dieback, are an expected risk of global trade of plant material^9,10^. In line with the propagule pressure theory^11^, *H. fraxineus* is a typical invasive species that relies on high ascospore production^12^.

The disease was first observed in Europe in north-western Poland in 1992 and, moving West, was identified in the UK in 2012^4,13,14^. The majority of ash trees are susceptible to the pathogen and less than 5% of trees are resistant or tolerant^15–17^. Ash dieback disease is characterised by dark brown/orange lesions on leaves followed by wilting, necrotic lesions on shoots then diamond-shaped lesions on the stems and finally preceding dieback of the crown^13,15,18^. The loss of leaves in the crown of mature trees proceeds over years and leads, in severe cases, to tree death. *H. fraxineus* is a heterothallic fungus^19^, shown to reproduce asexually *in vitro*^20^ but in the wild, probably through annual obligate sexual reproduction on fallen leaf rachises in the leaf litter^3,21^.

A pathogen's evolutionary potential, is rooted in its genetic diversity (i.e. its effective population size)^22^. Large populations can adapt more quickly than small ones for two reasons; first, a large population carries more polymorphism and so there is a greater chance that a favourable mutation is present in that population. Second, the impact of random genetic drift is lower for larger populations and therefore, natural selection is better able to drive those favourable mutations to fixation^23^. However, pathogen introductions are often associated with genetic bottlenecks which reduce the level of genetic diversity and the efficacy of selection^24^. This disparity between reduced polymorphism and invasion success is known as the genetic paradox of biological invasions^25^. The potato late blight pathogen, *Phytophthora infestans,* is one such example of a successful bottlenecked pathogen introduction^26^. However devastating a pathogen invasion may be, multiple introductions can increase diversity higher than that of native populations^27^. Early *P. infestans* invasions were dominated by a single clonal lineage (US-1) but this lineage was superseded by lineages of increasing diversity and increasing severity^28^. The Dutch elm disease pandemic(s) was also characterised by the initial intense spread of *Ophiostoma ulmi* which later declined and was replaced by O. novo-ulmi (also subsp. *americana),* which went on to kill elms across two continents^2^. The level of variation that has been introduced to Europe and the potential for further introduction are of critical concern and a key focus of the present work.

Estimates of *H. fraxineus* microsatellite allelic richness suggest as few as two haploid individuals may have invaded Europe^3^. Such a bottleneck would represent a genetic paradox and so it is important that we measure nucleotide diversity across the genome as well as meaningful, adaptive diversity in the host interaction genes (effectors). Furthermore, given the impact of the pathogen invasion so far, we set out to understand the characteristics of the source population and the signal of the introduction in order to highlight the consequences of further introduction of polymorphism via identifiable routes of intercontinental disease transmission. Here, we assembled and annotated a high-quality draft genome. We quantified the level of genetic variation in 43 *H. fraxineus* isolates from across Europe as well as 15 isolates from part of its native range (Japan). In order to understand the adaptive potential of the ash dieback pathogen in Europe today as well as its future potential through successive invasion, we determined the effective population size, quantified the bottleneck into Europe and the estimated the size of the source population. In addition, we measured signals of adaptive diversity in genes, including the secreted protein genes encoding for putative effectors. The proportion of introduced adaptive variation and the potential for further introduction are a key focus of the present work.

## Results & Discussion

Population genetic diversity across the genome of the emerging invasive *H. fraxineus* pathogen measured among 43 isolates from across Europe is an eighth that of 15 isolates from a single wood in Japan. This general reduction in polymorphism caused by a bottleneck of two divergent individuals could reflect a reduction in the pathogens adaptive potential in its introduced environment. This reduction in genetic diversity is present in effectors as well as all other genes. Effectors in Europe retain a signal adaptive variation but far less than that present in the native range and we discuss the implications for the predicted level of virulence in Europe over the long term.

### Hymenoscyphus fraxineus version2 genome approaches chromosome level contiguity

The *Hymenoscyphus fraxineus version 2* genome assembly (*Hf*-v2.0) is 62.08Mbp (Fig. 1). *Hf*-v2 contains the information present in 1672 contigs of *Hf*-v1.1 in just 23 scaffolds (Supplementary Information 2). Both *Hf*-v1.1 (published as a pre-print^29^ and *Hf*-v2 were released (open) online and hosted on the OADB github repository (https://github.com/ash-dieback-crowdsource^30^). The *Hf*-v2.0 gene annotation pipeline built on that of Saunders *et al.*^29^ and identified 10,945 genes (11,097 transcripts, 1516.50 mean CDS length, 3.49 mean exons per gene; Supplementary Information 3). Telomeric repeats are identified at both ends of 14 scaffolds and, at one end of all but one of the remaining scaffolds (Supplementary Table 2). Close linkage between three pairs of scaffolds missing a telomere was detected in the offspring of an *H. fraxineus* cross using SNPs at the ends of scaffolds. After joining these scaffolds, we had an estimate of 20 chromosomes (pseudomolecules) for *H. fraxineus* (Fig. 1 track 2).

**Figure 1.**
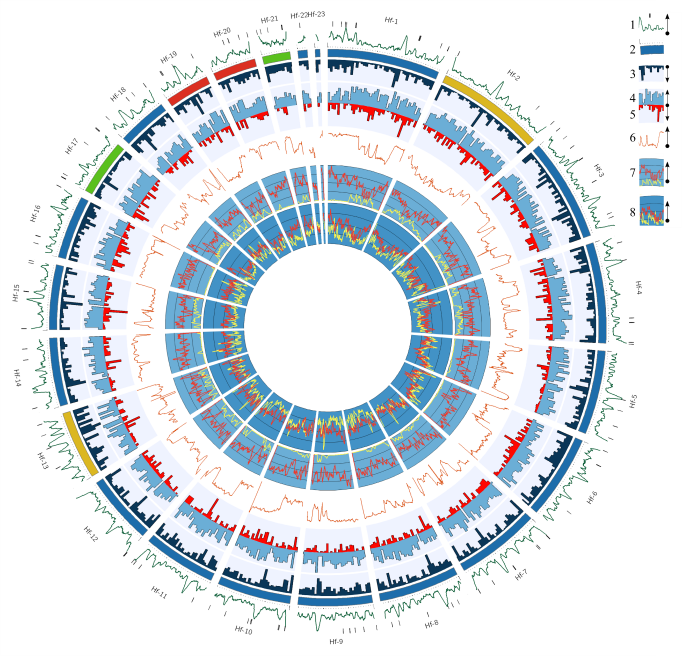
*H. fraxineus* genome organisation (repeats, effectors etc.) and population statistics (*F_ST_*, *π* etc.). (1) AT richness sliding window (63Kbp-5Kbp) with ticks indicating regions of n's or gaps in the assembly >2.5Kbp. (2) Scaffolds (ticks = 100Kbp) with evidence for linkage between red, green and yellow pairs of scaffolds. (3) Repeat density (100Kbp). (4 & 5) Gene density & effector density, respectively (100Kbp). (6) Genetic differentiation (*F_ST_*) sliding window between Japanese and European populations (100Kbp-5Kbp). (7) Nucleotide diversity *(π)* sliding window within Japanese (red) and European (yellow) populations (100Kbp-5Kbp). (8) SNP density sliding window within Japanese (red) and European (yellow) populations (100Kbp-5Kbp). Legend shows track number and orientation.

### The European invasion population is bottlenecked and sexually recombining diversity from Asia

In 43 European individuals and 15 Japanese, we identified 6.26 million SNPs overall (SNPs in Japan (Jp) = 4.5 × 10^6^; SNPs in Europe (EU) = 0.67 × 10^6^; Supplementary information 5). A network of all genes shows us that the European population is genetically divergent and bottlenecked from that of the native population (Fig. 2). The European population is significantly smaller than the Japanese in terms of genetic diversity as the level of nucleotide diversity (π) of 43 individuals in Europe is an eighth of that observed in just 15 individuals in Japan (Wilcoxon (π), Jp genome > EU genome: W(1257)=393210, p < 0.001; Fig. 2). This disparity in nucleotide diversity between Europe and Japan could, arguably, have been caused because the European population was founded from a population much smaller than that of Japan. Tajima's *D*^31^ is a statistic, centred around zero, that is sensitive to changes in effective population size (and/or the mode selection). In Japan, we observe a Tajima's *D* value close to zero across the genome (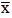 = −0.22) which indicates neutrality with purifying selection operating on genes which are more negative (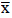 = −0.89; Fig. 2). In Europe, the genomic signal is broadly positive (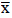 = 1.28; Fig. 2). This positive value is generated because the reduction in population size tends to remove high frequency alleles without equivalent reduction of rare alleles (Supplementary Fig. 10). This balancing of allele frequencies without a real reduction in SNP density by the loss of rare alleles across the genome is consistent with a bottlenecked founder EU population from a larger former population.

**Figure 2.**
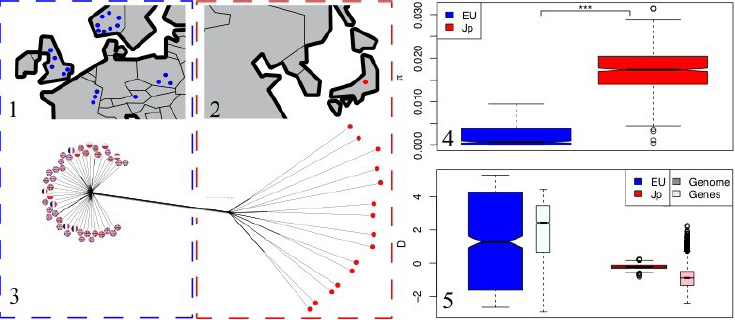
*H. fraxineus* European and Japanese sample sites and population genetic diversity. (1 & 2) European and Japanese sampling locations. (3) A network of the coding regions of all genes in the genome (16,589,637bp) bounded by dotted line indicating the sampling location in panels 1 & 2. (4) Nucleotide diversity *(π* 100Kbp windows) in both Japanese and European populations is greater in Japan than in Europe. (5) Tajima's D for all SNPs (100Kbp windows) across the genome and again in all genes within the Japanese and European populations. In Japan, a signal of neutrality is present across the genome and increases in the genes (purifying selection). The European population has a much broader positive distribution, consistent with a recent population bottleneck.

Sex is a fundamental determinant of plant pathogen adaptive potential not least because it allows recombination of host interaction genes, or effectors^32^. Here, as well as observing sexual reproduction between European isolates in the lab (Supplementary information 4) we see a breakdown in linkage disequilibrium in the wild in Europe (Supplementary Information 6). At the genome level Japanese and European populations are divergent *(Fst* = 0.58 (95% CI = 0.54 − 0.56)) but recombination decouples relationships between genes. Across the genome, genetic differentiation correlates positively with diversity present in the Japanese population (*R*^2^ = 0.18, p < 0.001) but negatively with diversity present in the European population (*R*^2^ = 0.51, p < 0.001). That is to say that the more diversity present at a locus in Japan the less likely it is to be shared with Europe and, the more diversity present at a locus in Europe the more likely it is to be shared with Japan; a signal consistent with directionality from east to west (Fig. 1, tracks 6 & 7, Supplementary Fig. 11). Continued gene flow from the native range into Europe is important to prevent because sexual reproduction between divergent demes may be important for plant pathogen adaptation^33^.

### The European population was founded by two individuals from a large diverse population

Burokiene *et al.*^21^ reported similar levels of genetic diversity (based on 11 microsatellite loci) between central European populations and those on the invasion front. The authors suggest that for introduced diversity to quickly spread to the invasion front either, the centre of genetic diversity quickly recombined and spread by range expansion, or only a small number of divergent *H. fraxineus* isolates arrived in Europe and present day diversity is the product of recombination among those divergent isolates. Here, we use third base positions of the coding regions of Core Eukaryotic Genes (CEGs) to distinguish between these scenarios and describe the invasion process. CEGs are highly conserved proteins essentially encoded in all eukaryotic genomes^34^. Their importance to fundamental eukaryotic biology makes the 387 CEGs we identified in all *H. fraxineus* individuals an ideal set to explore the European invasion whilst minimising the impact of processes associated with polymorphism generation operating on other genes^35^. The bottleneck into Europe has reduced the number of CEG haplotypes to 2.3 and 42% of all CEGs were monomorphic. Whereas, in Japan, the average number of haplotypes was 12.6 with no less than two haplotypes per locus. Strikingly, European CEGs are grouped into two divergent haplotypes and the majority of individuals have one or the other of the two main haplotypes. Furthermore, the level of divergence between those haplotypes reflects that across Japanese CEG networks (Fig. 3, Supplementary Fig. 18). It is our interpretation that these divergent haplotype pairs present in Europe have been introduction by two haploid founders as first suggested by Gross *et al.^3^.* Moreover, the divergence between these European haplotype pairs, without intermediates, represents the shadow of an ancestral population from which the European population was founded.

**Figure 3.**
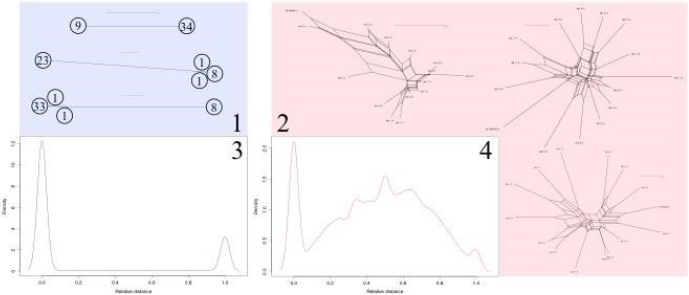
Divergence between haplotypes in Europe is high and bimodal. Networks were generated using the third base position of 387 CEGs. Three CEG networks are shown from European (1) and Japanese *H. fraxineus* (2) populations. Density plots show the relative pairwise distance amongst haplotypes of 387 CEG in Europe (3) and Japan (4). In japan, pairwise distances range in their divergence. In Europe, the majority of genes are either identical or at the complete opposite ends of the divergence distribution (numbered in circle). This represents the presence of two major divergent haplotype groups (Supplementary Fig. 18).

The observed estimate for the haploid effective population size (*&_π_* = *2N_e_μ*) in Japan is 2.46 million individuals (π = 0.0246) and the estimate from Europe is 0.67 million individuals (π = 0.00672; *μ* = 5×10”^9^ per base per generation). However, this European estimate of nucleotide diversity is inflated by the divergence shadow of the two haploid individuals that founded the population and we used this divergent polymorphism to estimate the size of the source population from which this divergence shadow was generated. Coalescent simulations of a hypothetical European source of fixed effective population size which was bottlenecked to two (haploid) individuals shows us that an increase in effective population size, increases the average divergence between haplotypes in the two bottlenecked individuals. We find that an effective population size between 2.2 and 2.8 million individuals (closest median estimate = 2.5M) most accurately replicates the observed diversity in present day European haplotypes (Fig. 4). Our estimate of the size of the source of the European *H. fraxineus* population is equivalent to that from a single Japanese wood. These estimates may reflect an expectation of the equilibrium diversity in any given *H. fraxineus* population but importantly also suggest that the European invaders could have come from a single site, perhaps even as a single ascocarp (fruiting body).

**Figure 4.**
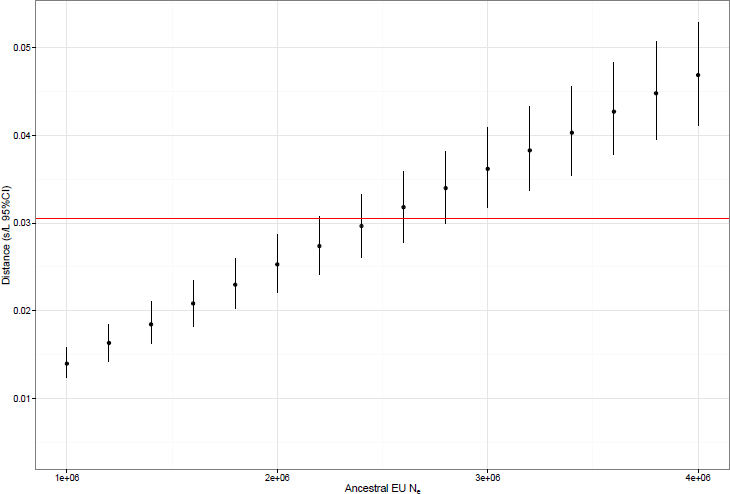
Coalescent simulations show a population of 2.5 million individuals maintain enough polymorphism to account for the observed haplotype divergence. The coalescent of a population was simulated (x 1000) at an effective population size of between 1M and 4M haploid individuals. At equilibrium, this population was bottlenecked to two individuals and the divergence at all loci with one or more polymorphisms was recorded. 2.5M individuals best encompassed the observed diversity in present day shared divergent haplotypes (observed SNPs/bp = 0.0305).

### 10% of H. fraxineus genes putatively interact with the host

The similarities between the effective population size of the Japanese and the estimated European-source populations suggest that the levels of genetic diversity observed in this native population reflect that of a representative European source. Effectors are a broad classification of proteins that interact with the host. They are secreted by bacteria, oomycetes, fungi, nematodes and aphids and they disable host defence components and facilitate colonisation. As such, effectors are in a co-evolutionary arms race with host resistance genes^36^ and therefore can be studied for insights into pathogen adaptive potential. We used localisation signals of N-terminal presequences^37^ to identify secreted proteins (i.e. putative effectors) throughout the genome of *H. fraxineus* (Fig. 1, track 5). From a starting set of 11,111 predicted proteins 1,132 are predicted effectors, which we clustered into 566 tribes (Supplementary Table 5). A search for N-terminal signal motifs found in putative effectors of other fungal pathogens did not reveal any conserved motif in *H. fraxineus* putative effectors. Density of AT base pairs are positively associated with repeat content (*R*^2^ = 0.79, p < 0.001; Fig. 1, tracks 1&3; Supplementary Fig. 19). Both genes (*R*^2^ = 0.68, p < 0.001; Fig. 1, track 4) and putative effectors (*R*^2^ = 0.08 p < 0.001; Fig. 1, track 5) are negatively correlated with repeat regions and putative effectors tend to be found in regions of increased gene content (*R*^2^ = 0.11 p < 0.001; Supplementary Fig. 20). Our observation that putative effector genes are spread across the genome similar to non-effector genes is consistent with the observation for the *Hf*-v1.1 genome^29^ and contrasts with that of the two-speed genome model, demonstrated in other filamentous plant pathogens^38^. Perhaps the requirement for a dynamic process that operates to shuffle effector genes and generate novel effectors is lower in the genome of a sexually reproducing pathogen with high haplotype diversity^35^. The presence of sex and high haplotype diversity in turn indicate a long term balanced relationship between host and pathogen within the native range^39^.

### Structure and function of H. fraxineus putative effectors

Of the 1094 predicted and manually curated *H. fraxineus* effectors, 64% carry Pfam signatures suggesting highly diverse biological function (Supplementary Table 6). The largest subgroups include 127 proteins predicted to have glycosyl hydrolase activity, 77 have oxidoreductase activity, 71 carry a cytochrome P450, 56 harbour conserved domains of unknown function and 396 did not carry a Pfam domain. To our knowledge, *H. fraxineus* is the first sequenced fungal pathogen with such high number of predicted Cytochrome P450 proteins, and most of them possess putative secretion signals. Cytochrome P450s mode of action is typically via the monooxygenase reaction which is important in the generation and destruction of chemicals especially aromatics. In *H. fraxineus* they may be important in pathogenesis, especially as they carry a signal of diversifying selection (Supplementary information 7). Potential roles for P450s in pathogenesis include: 1) destruction of ash tissue-derived aromatic compounds with antifungal properties. Previous studies have demonstrated that plant secondary metabolism pathways are active during the infection process of Magnaporthe oryzae^40^. 2) Penetration of ash tree tissues, P450 transcripts are upregulated in some pathogenic ascomycetes during the penetration of the plant cuticle, and are assumed to be used in the metabolism of hydrocarbon compounds which are the main constituents of host plants cuticle^41^. 3) Secreted *H. fraxineus* P450s could be used to facilitate fungal growth by extracellular oxidation of wood tissue, freeing nutrients, as well as enabling hyphal penetration^42,43^. 4) P450s are part of the biosynthetic pathways for mycotoxins and phytohormones secreted by *H. fraxineus* during invasion of ash tissue, similar modes of actions have been shown for Fusarium multifunctional cytochrome P450 monooxygenase Tri4^44,45^. Although most of the studied P450s in fungi are those with predicted intracellular activity, predicted secreted P450s in the ash dieback pathosystem could perform similar functions within specialised vesicles, the host apoplast or when transferred inside infected ash cells from invasive fungal structures. The presence of such high number of P450s predicted in the secretome of *H. fraxineus* suggests biochemistry unique to *H. fraxineus.* Thus, these P450s are potential targets for novel fungicides with selective inhibitory action, similar to azole inhibitors that are selective for the fungal P450 gene CYP51, critical in sterol biosynthesis^46^.

Transcribed low complexity domains may provide effectors with flexibility and diversity driven by adaptive evolution^47^. Of the 1094 effector set, 23% contain short repeats (e.g. Ankyrin repeats, Tetratricopeptide repeats and Leucine-rich repeats; Supplementary Table 7). Some are predicted to have a nuclear localisation signal and are indicative of putative intracellular function in host cells while others are part of chitin recognition domains such as LysM, revealing possible apoplastic function. A large proportion of effectors (38%), are cysteine-rich with one to five predicted counts of structural disulfide bonds (Supplementary Table 8) which are predicted to confer stability, biological activity and resistance to proteases in the apoplastic space^37^ and inside infected host cells^48^.

Based on Gene ontology (GO) and BlastP search against fungal and oomycete proteins, we identify three effectors (one of them previously identified^29^) with an NPP1 (necrosis-inducing Phytophthora protein) domain which is present in fungal, oomycete and bacterial proteins that induce hypersensitive reaction-like cell death upon infiltration into plant leaves^49^. Two other proteins have predicted cell death activity and yet two more carry S1/P1 nuclease activity domains, which are involved in non-specific cleavage of RNA and single stranded DNA^50^. Host cell death induction is suggested to be common among facultative parasites that engage in a necrotrophic lifestyle^51^.

### Adaptive effector diversity is present in Europe, but is far greater in the native range

In sexually reproducing species genes may carry their own demographic history and here we can use estimates of nucleotide diversity, SNP effects and non-synonymous to synonymous polymorphism to understand the roles of these genes. We analysed the level of nucleotide diversity in all the genes, (including putative effectors) of the European and Japanese populations (Supplementary Table 9). First, there remains a significant positive correlation in the level of genetic diversity present in genes between the founder European and Japanese populations (gene π, *R*^2^ = 0.180, p < 0.001; effector π, *R*^2^ = 0.144, p < 0.001; Fig. 5), despite a significant (73%) reduction in nucleotide diversity in Europe (Wilcoxon (*π*), Jp all genes > EU all genes: W(21885) = 102060000, p < 0.001; Fig. 6). This reduction in the level of nucleotide diversity has impacted genes (other than putative effectors) and putative effector genes alike as they are no different (Wilcoxon (π), EU genes ~ EU effectors: W(10942) = 5420600, p = 0.78; Fig. 6). Whereas, in Japan putative effector genes maintain significantly greater nucleotide diversity than that of other genes (Wilcoxon (π), Jp genes < Jp effectors: W(10942) = 5671200, p < 0.041; Fig. 6).

**Figure 5.**
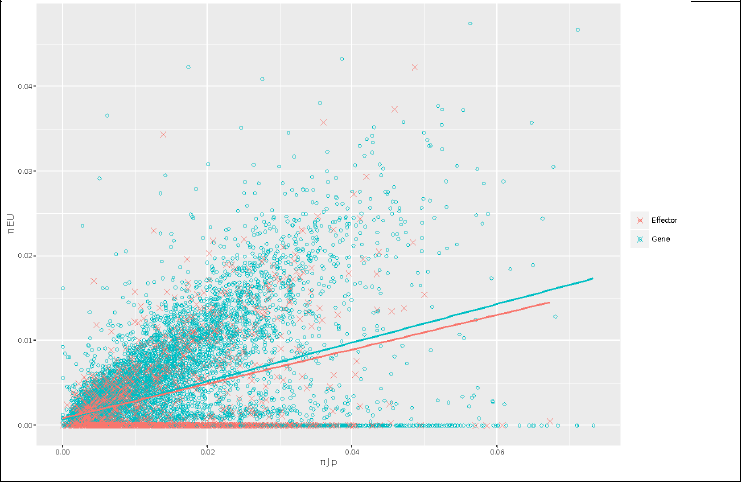
The nucleotide diversity (π) in all genes in the Japanese and European populations is positively correlated. This association demonstrates the association of polymorphism present in genes in Japan and Europe even after a population bottleneck. This reduction in nucleotide diversity for both genes and putative effectors in Europe is not significantly different between these groups of genes.

**Figure 6.**
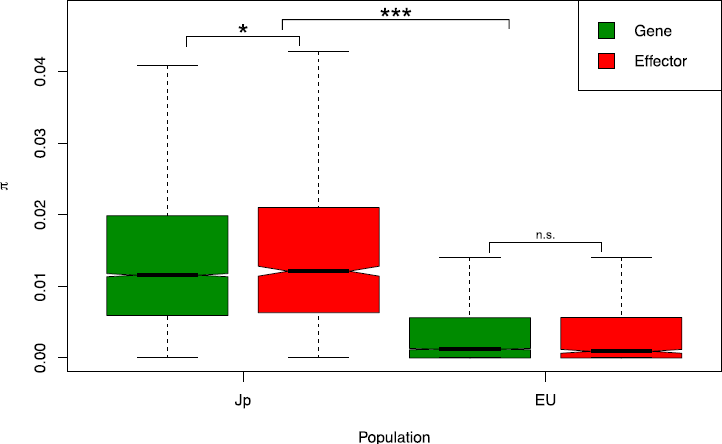
Nucleotide diversity (π) present at putative effectors and all other genes within Japanese and European populations. Nucleotide diversity (outliers not shown) over all genes combined is significantly greater in Japan (Jp) than in the Europe (EU). Effectors have significantly greater nucleotide diversity than all other genes in Japan but not in Europe.

SNP effects can be characterised and we find that putative effectors in Europe have an increase in SNPs found to affect splice donor sites, or found in splice regions, and 5' UTRs (Supplementary information 7). Importantly however, there are also increases in SNP densities in synonymous, intron, upstream and downstream positions in putative effectors relative to other genes (Supplementary information 7). An increase in SNP density can be the product of mutation (with neutrality) and linkage to a locus under balancing selection (Hughes & Yeager 1998). Therefore, observed increases within these effector features could be the product of increased SNP density through linkage and balancing selection operating on effectors, rather than an indication of the importance of the feature *per se.* The overall pattern of SNP effects was similar in both the Japanese and European populations despite the overall reduction in diversity in Europe (Supplementary information 7).

Pathogens are under constant pressure to avoid detection by their hosts (van Valen 1974). Pathogen effectors must adapt to avoid recognition by the host surveillance system and are expected to undergo positive, diversifying, and balancing selection^52,53^. To investigate the role of selection in the maintenance of polymorphism in putative effectors we assessed the level of adaptive diversity as measured using the mean pairwise ratio of non-synonymous *(ka)* to synonymous (ks) polymorphism present at all genes. The *ka:ks* ratio is useful to detect the presence of adaptive evolution when applied regions of genes, such as binding domains, because a value greater than one is consistent with positive or diversifying selection (functional divergence). Moreover, a *ka:ks* close to zero indicates negative or purifying selection. Here we apply the *ka:ks* ratio across whole genes, which, we presume have different modes of selection operating across them. However, we expect that effectors will retain an overall higher *ka:ks* value despite dilution by purifying selection operating over the majority of the rest of the gene.

Approximately 97% of all *ka:ks* values in the European and Japanese effectors are lower than one. Within this range, we observe a significant increase in the effector *ka:ks* over that of other genes between *ω* = 0.2 − 0.5 (Fig. 7). This represents a reduction in the strength of purifying selection in effectors relative to other genes. At *ka:ks* values greater than one, we observe a significant peak in the density for effectors which indicates a number of effectors evolving under contemporary positive selection (Fig. 7). Effectors in Japan have higher *ka:ks* values than other genes (Wilcoxon *(ka:ks),* Jp genes < Jp effectors: W(10942) = 4891800, p < 0.001; Fig. 8). In Europe, effector adaptive diversity has been maintained despite the bottleneck, albeit at a lower level than in the native range (EU genes < EU effectors: W(10942) = 1750600, p = 0.045; Fig. 8). Finally, consistent with linkage and balancing selection^54^, the level of synonymous polymorphism *(ks)* is also higher in effectors in both Japan and Europe (Wilcoxon *(ks),* Jp genes < Jp effectors: W(10942) = 4399800, p = 0.040; Wilcoxon (ks), EU genes < EU effectors: W(10942) = 1804900, p = 0.001; Fig. 8). We observe these *ka:ks* increases in effectors despite (presumed) purifying selection operating across a portion of the coding region. Pseudogenes too, may have a *ka:ks* ratio increased towards one. However, pseudogenisation could be more readily observed for effectors that are increasingly recognised by the immune system^55^. Here, we do not observe a significant increase in start lost, stop gained or stop lost variants in effectors over other genes (Supplementary information 7, Table 10) but more generally we believe that any impact of pseudogenisation on *ka:ks* is important to recognise.

**Figure 7.**
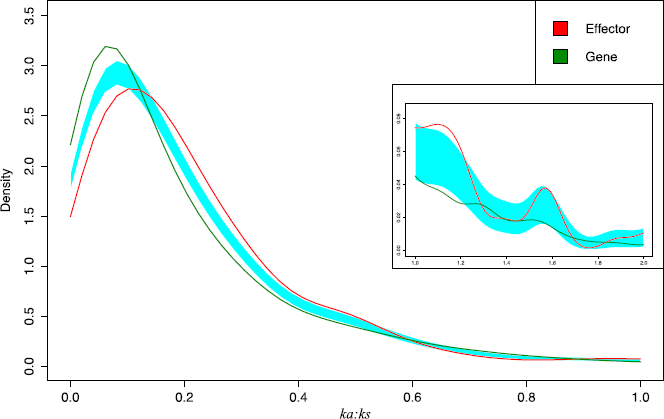
*ka:ks* density for genes and effectors in European and Japanese populations combined. *ka:ks* is significantly higher for effectors over a considerable range of the distribution between zero and one (p < 0.001). This is evidence for reduced efficacy of purifying selection in these genes. *ka:ks* greater than one (less than two; inset) also contains a peak for effector genes which is consistent with the operation of positive selection driving adaptive change in these effectors (p = 0.01).

**Figure 8.**
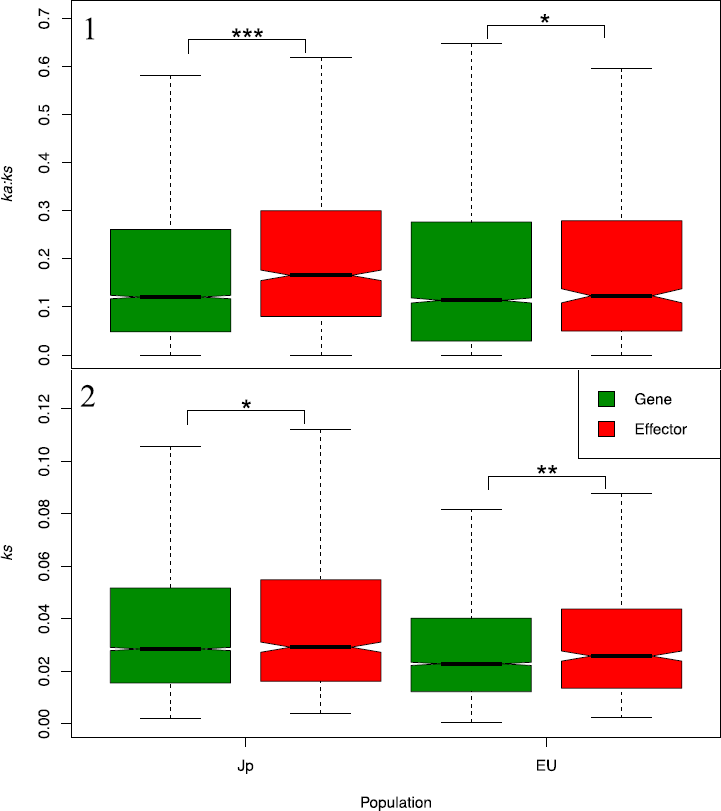
*ka:ks* and synonymous site density *(ks)* for effectors and all other genes. Outliers not shown. The signal of polymorphism visible to natural selection is greater in effectors than other genes in both Japan and Europe. However, the strength of this signal has been reduced in the European population. Synonymous site density is significantly higher in effectors in both the European and Japanese populations.

## Conclusions

The bottleneck of *H. fraxineus* into Europe removed the majority of neutral and much of the adaptive genetic variation. Despite this, the pathogen has decimated ash from East to West. Host defence is characterised in terms of levels of susceptibility^56^ and with native effector (and genome) diversity high, further introduction of pathogen genetic diversity runs the risk of increasing disease prevelence^57^. Successive invasions can both increase the level of genetic diversity above that of native populations and drive temporal fluctuations in important pathogen diversity^27,58^. Finally, an increased virulence at invasion onset, due to the density of uninfected hosts^59,60^, may be attenuated by natural selection on the pathogen as uninfected density reduces^61^ but this is much less probable where continued gene flow is allowed. For all of these reasons, it is most important to prevent further introduction of pathogen diversity to Europe.

Tree pathogens can be accidentally introduced because of commercial trade of live trees, wood and wood products^1^ and we are unable to manage host resistance in the same way that we do for crops^62^. If we interpret the divergence between founder haploids to mean that multiple native genotypes can invade, we face a situation in which the majority of genes (91.9%) from a single additional haploid individual from the Japanese population would harbour a novel allele. Given the overall number of haplotypes present in Europe and Japan, any further migration from the East is likely to be accompanied by extreme increases to the level of adaptive polymorphism and disease severity of the European population.

Here we publish a high-quality *H. fraxineus* genome with annotation and we begin to analyse the population genetics of the European invasion using a limited sample of the native range. Further population genetics analyses including broader sampling of single isolates from the native range will allow robust modelling of the source meta-population and the invasion scenario. Moreover, population genetics of *de novo* assembled isolates can be used to unravel processes of genome evolution given the specifics of this two-haploid introduction. Important for the understanding of the disease progression is the combination of native effector nucleotide statistics with targeted RNA-seq experiments.

Notwithstanding the potential impact of an introduction of the emerald ash borer^18,63^, current levels of European pathogen genetic diversity may infect 95% of all European ash. We must consider the implications of further introduction of diversity from the ash dieback pathogen.

## Materials & Methods

### Hymenoscyphus fraxineus collection and sequencing

A total of 44 *Hymenoscyphus fraxineus* isolates from Europe (21 UK, 13 Norway, 5 France, 4 Poland, 1 Austria) and nine from Japan (one haploid and eight fruiting bodies) were collected and cultured on malt extract agar^13^. Species confirmation was performed by assignment using PCR and sequencing of ITS sequences. DNA was sequenced at either Edinburgh Genomics or The Earlham Institute on MiSeq and HiSeq Illumina platforms to between 25-160 sequence depth (Supplementary Table 1).

### Genome Assembly and annotation of H. fraxineus

As part of a commitment to open science and rapid community analyses, the genome of isolate KW1, *Hf-*v1.1, was released online in March 2013 on the open ash dieback (oadb.tsl.ac.uk) repository^29^. *Hf-v2,* published here and released online in February 2015, was assembled using a 200bp insert Paired End (PE) and a Long (5Kbp) Mate Pair (LMP) library. Overlapping PE reads were merged together using FLASH^64^ and LMP reads were processed using FLASH and NextClip^65^ with modifications as described in^66^. SOAPEC error correction (k=25, low coverage cut-off: PE = 4, LMP = 7) and SOAPDENOVO v2.04^67^ (pregraph −K 111 −p 200 −R; contig −M 1 −R −p 200; map; scaff) were used to assemble and scaffold (Supplementary Information 2).

The annotation pipeline used for genome Hf-v1.1^29^ was replicated on Hf-v2 adding newer RNA-seq data available via the Open Ash Dieback repository (Supplementary Information 3). The *Hf*-v2 genome was repeat masked using *de novo* modelled repeats, known fungal and low complexity repeats (REPEATMODELER v1.0.8, REPEATMASKER v4.0.5^68^). Annotations (9,737) from Hf-v1.1 were transferred to Hf-v2 using GMAP^69^. AUGUSTUS v2.7^70^ was used to predict additional protein-coding genes using protein and RNA-seq alignments (16 libraries , Supplementary Table 16) with the previously trained *Hymenoscyphus fraxineus* model. We ran a transcript extension protocol on all genes to identify whether those genes within 100bp of an upstream gene had evidence for extension. In addition, we checked this extended gene dataset for potential secreted proteins. This protocol allowed the identification of 36 genes that had not previously been recognised as having a methionine start codon which were included into our effector-mining pipeline.

We used the pipeline described by Saunders et al.^29,37^ to identify putative effectors. Briefly, SignalP4 (−t euk −s notm −u 0.34) was run to identify transcripts with a signal peptide and from these TMHMM and TargetP were used to remove transcripts with transmembrane or mitochondrial localisation signals^71^. Finally, secreted proteins were clustered into tribes using TribeMCL vl2.135 (e-value 10^-5^)^72^. We used T-REKS^73^ to identify repeats, the Disulfide algorithm^74^ to predict disulfide bonds and PREDICTNLS^75^ to predict nuclear localisation signals.

### Mapping and SNP calling

Reads were trimmed for Illumina adapters and Phred quality (-q20) and discarded where length was reduced below 80bp (Trim Galore v0.3.3; Babraham Institute, Cambridgeshire, UK). Read alignment and mapping were performed using BWA-MEM v0.7.7^76^. SAMtools v0.1.19 and BCFtools v0.1.19-44428cd^77^ were used to sort, remove duplicate reads, mpileup (-D) and call variants (bcftools view −cg). VCFtools v0.1.13^78^ was used to filter variants to a minimum depth of ten, maximum depth of 1.8 × mean individual depth and a SNP genotype quality of at least 30. SNP sites with more than two alleles were excluded as likely errors, as were sites reported as heterozygous in haploids. Finally, sites that were missing in 20% or more individuals were also removed. Three samples were removed from further analyses because of insufficient depth or evidence of contamination (20 UK, 13 Norway, 5 France, 4 Poland, 1 Austria, 8 Japan (1 haploid 7 fruiting bodies); see Supplementary Information 1).

### Phasing fruiting body samples

SHAPEIT2^79^, used to phase fruiting body data, first applies a phase informative read step (extractPIRs) to group SNPs present on the same read pair and then phases remaining SNPs using population level polymorphism in the second step (assemble --states 1000 --burn 60 --prune 60 --main 300 --effective-size 564000 --window 0.5). All Japanese samples including the haploid strain were used in the phasing step. As part of the phasing process indels must be removed from the dataset. Effective population size (*θ=2N_e_μ*) was calculated assuming a mutation rate of 5 × 10^-9^ per generation per site^80^ using the mean nucleotide diversity present in 100Kbp windows in the Japanese population.

### SNP diversity and divergence analysis

Polymorphism statistics and sliding windows i.e. sites per individual, SNP depth, transition/transversion (Ts/Tv) ratio, linkage (--thin 1000; --ld-window-bp 500000), *π* and *Fst* were conducted using VCFtools v0.1.13 & BEDTools v2.22.0^81^ & custom scripts (https://github.com/mcmullan0). SnpEFF v4.2^82^ was used to categorise the impact of variants. Effectors are, on average, shorter than other genes and so we controlled for the number of variants present per feature by dividing by feature length. DNAsp v5.10.01^83^ was used to calculate population genetic statistics per gene after fasta conversion using GATK v3.5.0 FastaAlternateReferenceMaker^84^ and PAML v4.9^85^ was used (YN00) in pairwise mode to calculate the average *ka:ks* ratio nonsynonymous (ka) and synonymous (ks) values for each gene with at least one synonymous mutation. The sm package^86^ was used in R v3.2.1 to compare the density distributions of *ka:ks.* Output data for each gene (including effectors) from SnpEff, DNAsp & PAML are combined into a single table. Data are represented using SplitsTree v4.13.1^87^ using default parameters (CDS of all genes and separately for each CEG) and CIRCOS v0.67-5^88^.

### Parental cross and linkage analysis

Two *H. fraxineus* isolates of opposite mating types, LWD054 and LWD067, were collected from stem lesions on trees in Norfolk, UK in 2014. Crosses were made between parental lines according to the method of Wey *et* al.^89^ except that isolates were kept at 25° C in 16 hours light. KASP™^90^ markers were designed based on SNP sites between two *H. fraxineus* parents mapped to the *Hf*-v2 assembly (Supplementary Information 4). Linkage between scaffolds was ascertained using Chi-squared test for linkage of sites close to ends of the 23 scaffolds, across 28 F1 offspring. Scaffolds at linkage group ends were examined for the presence of telomeric motifs.

### Statistics

R v3.2.1 was used to fit correlations to test the level of association between population signals in order to show directionality in the invasion and portray the bottleneck behind Tajima's D. In addition, we correlated feature coverage (i.e. repeat content) of regions with gene and effector coverage to understand genome evolution. Chi-squared tests were used to assess linkage between scaffolds in the offspring of a parental cross. Two-sample Wilcoxon tests (one tailed indicated by '>' or '<'; two tailed indicated by '~'), were used to test for differences in signals of nucleotide diversity, Tajima's D and *ka:ks* between non-effector genes and effectors in European and Japanese populations. Bootstrap analyses were used to compare variants effect by feature or gene (e.g. SnpEff output) and involved sampling feature with replacement (×1000). Bootstrapped comparisons are tested using a randomisation test and 95% confidence intervals (CI) are presented.

### Ancestral effective population size

In order to estimate the size of the population from which the European population was founded we used the SNPs within the third base pair of codons within 387 core eukaryotic genes. CEGMA v2.4^34^ initially identified 440 CEGs which was reduced to 387 genes that were confirmed by reciprocal blastp (BLAST v2.2.31^91^ (-evalue 1e-5) with a minimum of 90% coverage (to remove spurious technical errors in the assembly and/or biological duplicates), removal of CEGs with nonsense variants (expected pseudogenes) and a minimum mean depth of 10x per gene across all individuals (remove poorly covered CEGs). Each gene was then run through a pipeline to quantify the pairwise distance between each allele (using SPLITSTREE) and the diversity per gene (using DNASP V5).

The average number of base pairs (3^rd^ codon bp) per CEG was 448 and this number was used as an input to fastsimcoal2 v2.5.2.8^92^, a fast sequential markov coalescent simulator used to estimate the size of the population from which the founders into Europe came. A simple model (x1000) of a single (haploid) population of fixed effective size (*N_e_* = 100,000 − 4,000,000) containing 387 unlinked CEGs that were freely mutating at a rate of 5 × 10^-9^ per base per generation^80^ was bottlenecked to two individuals. From these two individuals, we recorded the number of haplotypes and the haplotype divergence at each gene for comparison to observed data.

## Data access

All MP and LMP sequencing data generated for the Hfv2.0 genome (PRJEB21027), population genetics and parental cross reads have been submitted to the European Nucleotide Archive (under projects PRJEB21061, PRJEB21062, PRJEB21063, PRJEB21064, PRJEB21059, PRJEB21060; see Supplementary Table 1 for accession 's). The genome annotation is available at the Earlham Institute Open Data site (http://opendata.earlham.ac.uk/Hymenoscyphus_fraxineus/EI/v2/).

## Author contributions

MM, CF, JB, MB, AD & MC contributed to the design of the project. EO, AE, KY, TH, LW, PJ, RI, CH, AH, AV, HS & AD contributed to collection of samples. Acquisition and processing of sequence data was done by MM, WV, AE, JW, GeoK & DM. MM, MR, GemK, BC, GC, LB, EO, DGS, LPA, BW, DS, JB, & MC contributed to the analysis and interpretation of data. MM, MR, GK, BC, LPA, LB, EO, AH, NH DS JB AD & MC contributed to the drafting and revising of the MS.

## Acknowledgments

We thank The Sainsbury Laboratory for hosting the Open Ash Dieback Git repository (oadb.tsl.ac.uk), and Sophie Janacek for assistance with project management. The submission of sequencing data was brokered by COPO (http://copo-project.org/). The Ash Dieback research project was funded jointly by the Biotechnology and Biological Sciences Research Council (BBSRC) and the Department of Environment Food and Rural Affairs (Defra) via the 'Nornex' project (BBS/E/J/000CA523) and Economic and Social Research Council, the Forestry Commission, the Natural Environment Research Council (NERC) and the Scottish Government, under the Tree Health and Plant Biosecurity Initiative (BB/L01291X/1). The UMR IAM is supported by a grant overseen by the French National Research Agency (ANR) as part of the “Investissements d'Avenir” program (ANR-11-LABX-0002-01, Laboratory of Excellence ARBRE). Genome sequencing was done by Edinburgh Genomics, and at the Earlham Institute (EI, formerly The Genome Analysis Centre, Norwich), by members of the Platforms and Pipelines Group funded by BBSRC National Capability in Genomics grant (BB/J010375/1). Additional research funding was via a BBSRC Institute Strategic Programme Grant for Bioinformatics (BB/J004669/1).

